# Neural basis of self-control

**DOI:** 10.1101/2024.02.07.578652

**Authors:** Ka Eun Lee, Jacob Elsey, Jaewon Hwang, Erik E. Emeric, Veit Stuphorn

## Abstract

Self-control is the capacity to inhibit self-defeating behavior in the face of temptation. The neural basis of self-control remains elusive, due to the difficulty of disentangling self-control from value-based economic choice. To overcome this problem, we designed a novel task for monkeys that allows to identify states of high and low self-control. We found that neurons in the Supplementary Eye Field encode self-control and predict whether and when monkeys give in to temptation. This neuronal activity was present early in the trial, even preceding presentation of choice options. Our findings suggest that Supplementary Eye Field is part of a neuronal circuit exerting proactive self-control, which is crucial for selecting and maintaining the pursuit of costly goals that are beneficial in the long run.

**One-Sentence Summary:** Supplementary eye field encodes proactive self-control signals independent from cost-benefit estimation.

---

Long-term goals require self-control, which involves foregoing tempting but suboptimal rewards to achieve larger, but more costly rewards. Despite the importance of self-control in everyday life, its neuronal mechanisms remain elusive. Traditionally, self-control has been studied using the intertemporal choice task (*1–9*), where the choice of a delayed larger reward (L) over an immediate smaller reward (S) is viewed as high self-control. However, such choices can also be the result of a cost-benefit evaluation of the offered rewards (*10–12*). For instance, choosing S over an L with a long delay can be rational if the subjective cost of the delay outweighs the value difference. Thus, the influence of self-control and value-based evaluations are conflated in intertemporal choice, making it challenging to isolate the neuronal mechanisms unique to self-control. To uncouple the two processes, we designed a modified version of the intertemporal choice task which allows to identify different levels of self-control independent of value-based choice (**Fig. 1A**). We recorded from the Supplementary Eye Field (SEF) (**Fig. 1C**) (*13*), to identify neuronal activity associated with self-control.

**Fig. 1.**
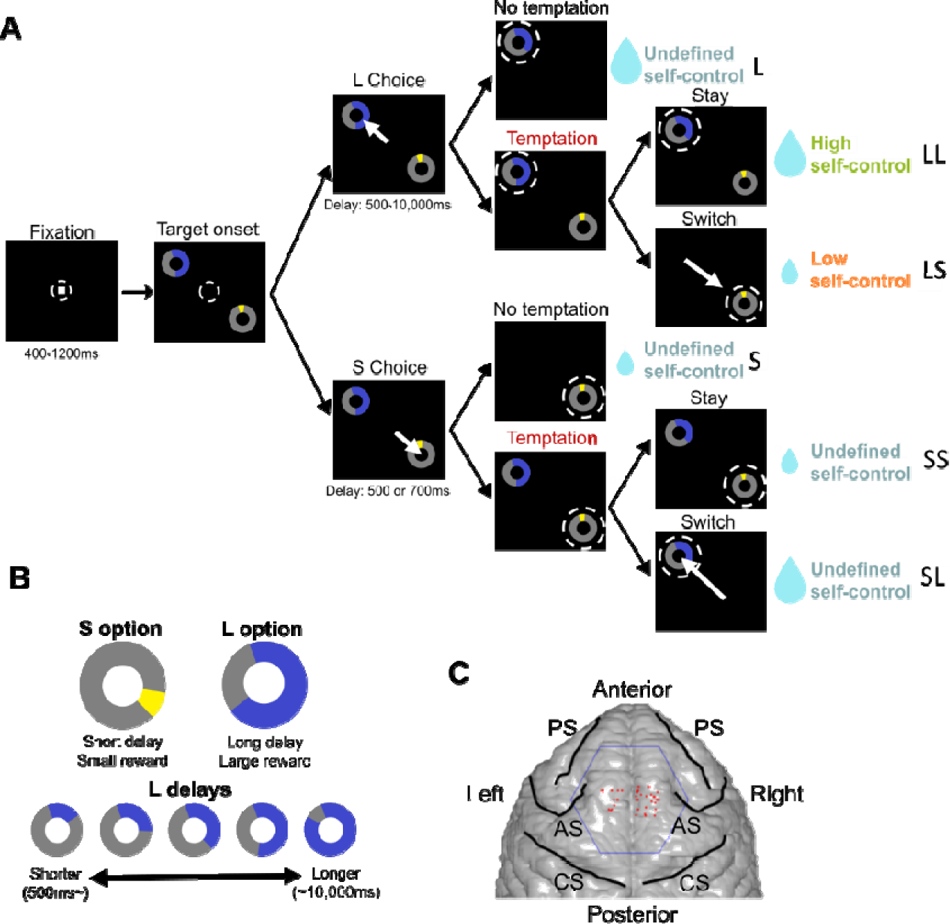
Self-control task design and recording locations. (**A**) Task design. After target presentation, the monkey chooses between S and L options by making a saccade to an option and remaining fixated until the end of the delay period. Dotted circles indicate the location of fixation by the monkey. White arrows indicate eye movements of the monkey. On temptation trials, the unchosen option remains available to allow switching from the initial choice. The final decision to switch from, or stay with, the initially chosen option is referred to as final choice. (**B**) Choice targets. The delay to reward is indicated by the area of space colored within the circular cue, and reward amount is indicated by the color. S delay was fixed within each session. L delay varied from trial to trial, ranging between 500ms and 10,000ms. (**C**) Example recording sites for a monkey. PS: principal sulcus, AS: arcuate sulcus, CS: central sulcus.

## Identifying self-control independent of cost-benefit estimation

Five rhesus monkeys (*Macaca mulatta*) performed the self-control task (**Fig. 1A**). On each trial, monkeys initially chose between a smaller sooner reward (S) and a larger later reward (L), which had explicitly cued reward amounts and delays (**Fig. 1B**). The L option was always the prudent option that maximizes reward rate (see **Materials and Methods**).

In the majority of trials (‘no-temptation’ trials), the unchosen option disappeared after initial choice, as in a typical intertemporal choice task. In contrast, in ‘temptation’ trials, which occurred less frequently (mean across sessions: 36.5% of trials; range: 10-54%), the unchosen option remained available after the initial choice. This allowed the monkey to potentially switch to the initially unchosen option. Whether the monkeys switched or stayed determined their final choice of S or L.

Because temptation trials occurred unpredictably, monkeys were unable to anticipate whether the unchosen alternative will remain available. Consequently, initial choices for no-temptation and temptation trials were not expected to be different. Instead, they should reflect a common cost-benefit evaluation. Thus, an initial choice of the L option indicates that on that trial, the larger delayed L reward, despite its longer delay, was deemed subjectively superior to the S reward.

Since the initial choice reflects the subjectively preferred option on all trials, switching immediately afterwards would be irrational from an economic point of view. What then could motivate switching? The L reward is not only associated with a higher reward, but also higher costs, creating a temptation to avoid them by switching to the immediate reward. The behavioral response to this temptation, when the monkey has chosen the L option, reveals therefore the level of self-control on that trial: maintaining the initially chosen L reward despite the costs (LL trials) indicates high self-control, while switching to the S reward (LS trials) to avoid the costs indicates low self-control. In this way, the same value-based decision (i.e., initially choosing L) can be made with different levels of self-control, allowing us to disentangle signals associated with self-control and value-based choice.

There are only two economically rational alternative explanations for switches. One is error correction (i.e., correcting an initially mistaken choice). However, error correction can be distinguished from self-control because it predicts symmetrical switches (i.e., L to S and S to L switches occur equally often), while low self-control predicts asymmetrical switches (i.e., switching predominantly occurs from L to S). We address this possibility in the following section.

The second is a dynamic readjustment of the L and S values in the waiting period following the initial choice (*14–16*). According to this hypothesis, the subjective value estimate of both options is continuously adjusted as both rewards get closer to being available. Because temporal discounting is hyperbolic, the subjective value estimations of L and S options can reverse, leading to a switch. We tested this idea by adding a 1-s additional delay to both options. This should have led to later switch times. However, we found that switch times stayed unchanged, arguing strongly against temporal discounting as the cause of switches (**fig. S1**).

Self-control level is unknown for initial S choices, because the effect of self-control and value-based preferences on choice cannot be distinguished (see **Materials and Methods**).

## Effect of temptation on maintaining prudent choices

To quantify the effect of temptation on choice behavior, we performed logistic regression on the choice data and computed the indifference delay (i.e., the L delay at which the monkey is indifferent between L and S).

All five monkeys made initial choices according to the delay-discounted subjective values of options (**Fig. 2A,B**). As predicted, the choice functions largely overlapped (**Fig. 2B**). Although we did observe a small, but significant difference (paired t-test: t=3.931, p=1.27e-04) (**Fig. 2E**), this was an artefact of the analysis design and not indicative of genuine anticipation (**fig. S2**).

**Fig. 2.**
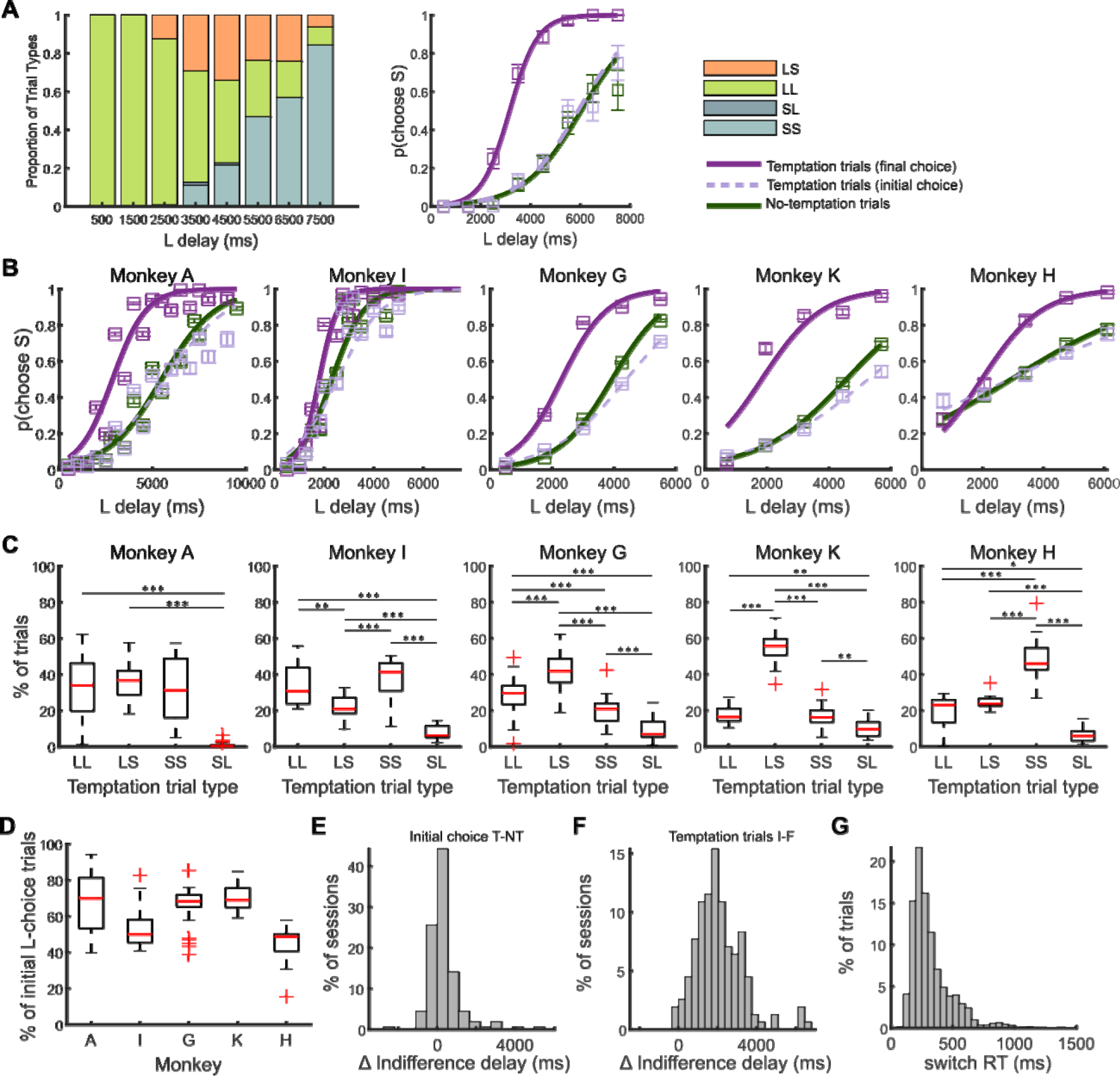
Behavioral results. (**A**) Proportion of trial types (left) and choice function (right) for an example session. (**B**) The proportion of trials where the S option was chosen as a function of L delay. Two different S, L reward combinations were interleaved within each session for monkey K. Only the trials from the larger reward combination were used for visualization. (**C**) Proportion of temptation trial types. Difference in trial type proportions was statistically significant in all four monkeys (one-way ANOVA, monkey A: F=82.25, p=8.54e-34; monkey G: F=103.75, p=3.27e-36; monkey I: F=32.73, p=6.07e-13; monkey K: F=233, p=1.34e-23; monkey H: F=36.29, p=5.58e-11). Stars represent the result of multiple comparison of group means (*: p < 0.05, **: p < 0.01, ***: p < 0.001). (**D**) Proportion of initial L choices. (**E**) Distribution of the difference in indifference delay for initial choice in temptation (T) and no-temptation (NT) trials. (**F**) Distribution of the difference in indifference delay for initial choice (I) and final choice (T) in temptation trials. (**G**) Distribution of switch response time (switch RT). Switch RT indicates the time from choice until switch, in no-temptation switch trials (LS, SL).

The final choice on temptation trials indicated a much greater preference for shorter delays than in the initial choice (mean difference in indifference delay_temp initial-final_: 2,110ms; 48.5% of initial indifference delay; paired t-test: t=11.04, p=2.87e-21). This shift reflects the asymmetrical influence of temptation on the monkeys’ preferences, resulting in more LS (86.3%) than SL (13.7%) switches (one-sample Wilcoxon signed-rank test against 50%; W=9037, p=5.76e-24) (**Fig. 2C**), indicating a strong bias toward selecting the S option under temptation. This imbalance demonstrates that most switches were not due to error correction. Error correction should cause LS and SL switches to occur equally often, resulting in a steeper choice function but no net change in preference. We did not observe such an effect (**Fig. 2B**). Moreover, this asymmetry was not the result of a physical incapability of the monkeys to switch from S to L, as 85% of all switches occurred before 500ms (median=276ms), the shortest S delay used in the task (**Fig. 2G**), nor was it caused by difficulties of the monkeys to remain fixated on the L option for the length of its delay (**fig. S3**). Thus, temptation prompted monkeys to forego the prudent choice and settle with the immediate reward, revealing trials with low self-control.

## SEF neurons encode distinct levels of self-control

The monkeys have variable success in maintaining prudent choices on temptation trials. To examine whether SEF neurons carry a signal that can influence the monkey’s response to temptation, we recorded from 822 SEF neurons in four monkeys performing the self-control task.

We first tested whether SEF neurons carry self-control signals by using generalized linear models (GLM) (see **Materials and Methods**). We examined SEF activity aligned to three events (fixation onset, target onset, initial choice) in overlapping time bins (75ms shifted by 25ms) (**Fig. 3A**). The variables and interactions tested are listed in **table S1**. For each time bin, we found the set of variables that best explained each neuron’s firing activity. A neuron was considered to encode a variable in a given bin only if it was included in the best-fitting model for 4 consecutive bins (i.e., present for at least 150ms, accounting for bin overlap) (**fig. S4**). Because we found a small but significant difference in choice reaction time (choice RT) for high (LL) and low (LS) self-control trials (**fig. S5**), we included choice RT as a potential variable to dissociate the effects of self-control and motor preparation on explaining neuronal activity.

**Fig. 3.**
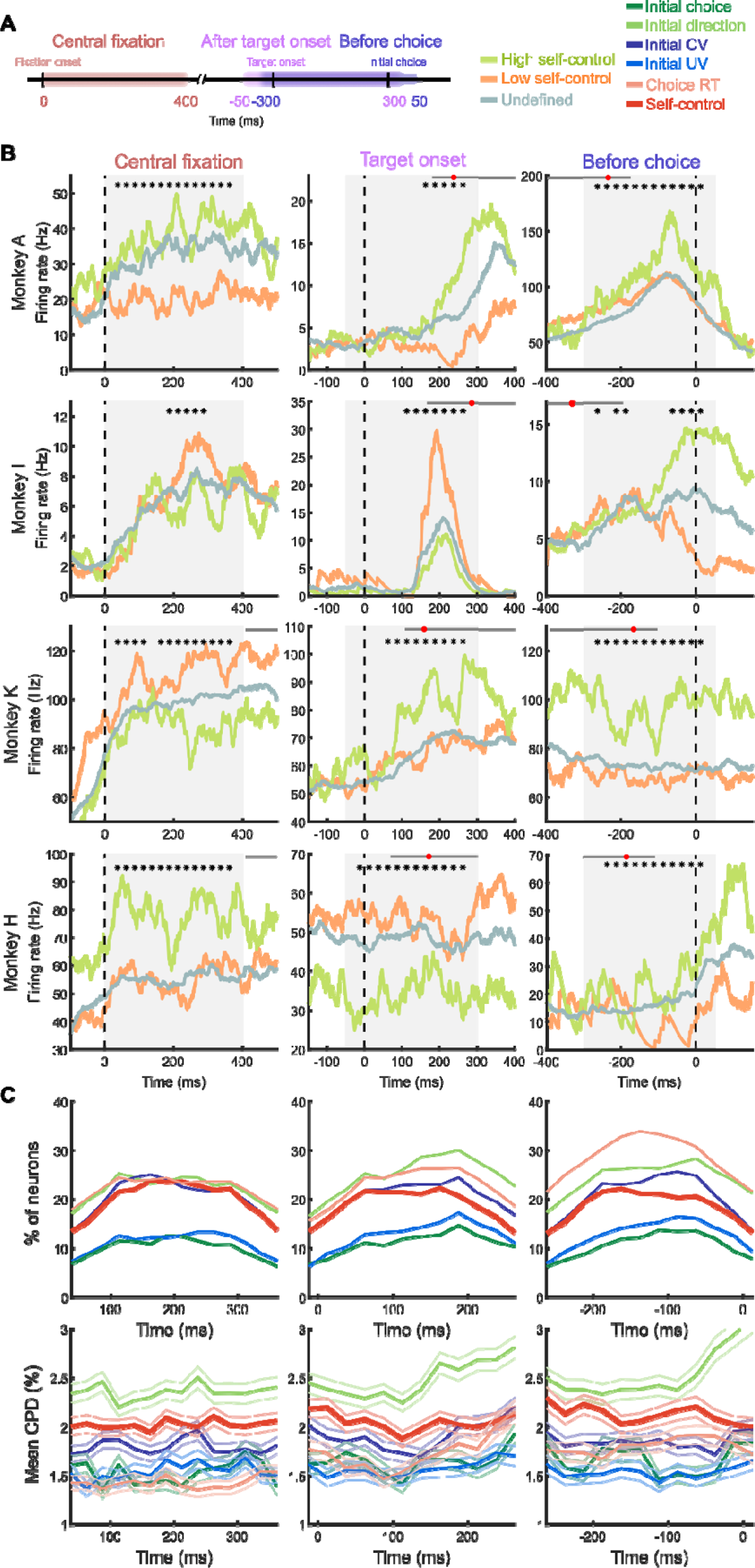
Encoding of self-control in SEF neurons. (**A**) Analyzed time periods and their alignments. Within each time period, SEF activity was analyzed in 75-ms time bins shifted by 25ms. Neuronal activity for central fixation, target onset and before choice was aligned to the time of fixation, target onset, and initial choice, respectively. (**B**) Spike density functions of example single neurons that encode different levels of self-control. Each panel is a different neuron. Stars indicate time bins when the neurons encoded self-control. Black horizontal bars for central fixation, target onset and before choice indicate the range of target onset, time of choice and target onset, respectively, in trials where the given neuron was active. Red dots indicate the median. Gray area indicates time period analyzed. (**C**) (top) Proportion of neurons encoding each variable in a given time bin. (bottom) Mean coefficient of partial determination (CPD) for each variable in each time bin. The shaded areas represent SEM.

SEF neurons encoded self-control in all 3 time periods (**Fig. 3**; see **fig. S6** for additional example neurons). Neuronal activity could be either positively or negatively correlated with high self-control (**table S2**). At fixation (0 to 400ms around fixation), 36.6% of neurons (301/822) encoded self-control. On average, 19.9% of neurons (mean=163, SD=30.5 neurons) were encoding self-control at any given time bin during fixation (**Fig. 3C**, **fig. S7**). Because choice targets were presented at the earliest 406ms after fixation onset (mean=872ms, SD=236ms), self-control signals in this time period were unlikely to be related to saccade preparation.

We next examined SEF activity in the time period after target onset and before initial choice (mean choice RT=228ms, median=214ms, SD=72.3ms). In this time period, the S and L targets are presented, starting the decision process culminating in the initial choice. Because SEF activity could reflect both sensory responses to the targets or cognitive and motor-related processes involved in choice, we analyzed the activity both aligned on target onset and initial choice. Aligned to target onset (−50 to 300ms), activity in 263/822 neurons (32%) encoded self-control, 167 of which (63.5%) had also been identified to encode self-control during fixation. On average, 18.8% of neurons (mean=154, SD=26.8 neurons) were encoding self-control after target onset at any given time bin (**Fig. 3C**). Aligned to initial choice (−300 to 50ms), activity in 264/822 neurons (32.1%) encoded self-control, 165 of which (62.5%) had also been identified to encode self-control during fixation, and 182 (68.9%) after target onset. On average, 18.6% of neurons (mean=153, SD=26.6 neurons) were encoding self-control at any given time bin (**Fig. 3C**).

Self-control was encoded by a consistent proportion of neurons in all three time periods. Overall, 443/822 neurons (53.9%) encoded self-control in at least one of the three time periods. 129/822 neurons (15.7%) encoded self-control in all three time periods (**table S3**). In addition to self-control, a substantial proportion of SEF neurons also encoded other choice-related variables, such as the spatial location and the delay-discounted subjective value of the initially chosen option (**Fig. 3C, table S4**). Some neurons encoded choice-related variables in the fixation period (**Fig 3C**), which was unexpected, because no choice-relevant information was present in that period. This might reflect self-control indirectly (see **fig. S8**).

Many neurons that encoded self-control also encoded interactions between self-control and other variables (neurons encoding self-control with an interaction with any variable: fixation: 168/301 (55.8%), target onset: 161/263 (61.2%), initial choice: 154/264 (58.3%); **table S5**). Most instances of interactions were found with *initial chosen value* and *initial direction* (**table S4**). Thus, self-control likely modulates the representation of variables such as the spatial location or subjective value of the options to influence initial choice and saccade vigor.

## Fluctuations in SEF activity predict monkeys’ response to temptation

The preceding analysis demonstrated that the activity of SEF neurons is correlated with different levels of self-control. Since these signals were present before initial and final choices, they are in a position to directly influence the monkey’s response to temptation. We therefore examined how well the final choice of the monkey can be predicted by trial-to-trial variations in the neuronal activity of SEF neurons. Only temptation trials were used for this analysis.

We collected neuronal activity in overlapping 75-ms time bins shifted by 25ms in two time periods: (1) −200ms to 300ms around target onset, and (2) −200ms to 300ms around initial choice (**Fig. 4A**). We searched for neurons whose activity explained variance in final choice over and above other choice-relevant variables for at least 4 consecutive time bins.

**Fig. 4.**
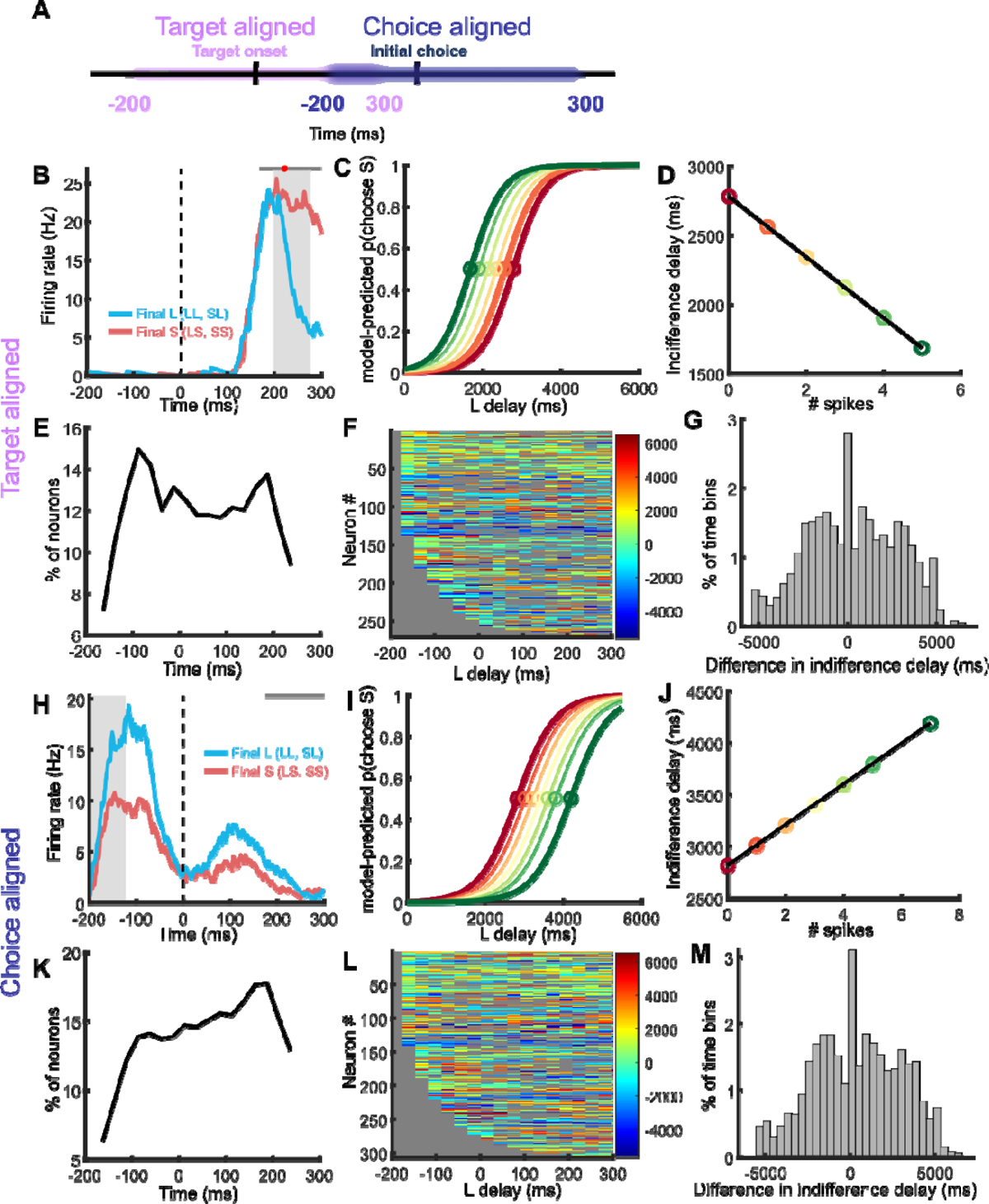
SEF neurons predict final choice in response to temptation. (**A**) Analyzed time periods and their alignments. (**B**) Spike density function of an example SEF neuron. Black bar indicates the range and red dot indicates the median of choice RT. Gray area indicates the time bin from which panels **C** and **D** were created. (**C**) Set of choice functions, each predicted by a given activity level observed from the neuron in the 75-ms time bin. Red and green choice functions indicate the prediction for the minimum and maximum firing rate, respectively. (**D**) Model-predicted indifference delays as a function of spikes. (**E**) Proportion of SEF neurons whose activity predicted final choice when aligned to target onset, in each time bin. (**F**) The difference in the estimated indifference delay for maximum and minimum firing rates of each given neuron, plotted over time for all neurons that predicted final choice, sorted by the onset of predictive activity. Color bar: difference in indifference delay (ms). (**G**) Distribution of the difference in estimated indifference delay_max-min_ for maximum and minimum firing rates of neurons that predict final choice. (**H-M**) Same as in **B-G**, but for the choice-aligned period.

We found that trial-to-trial fluctuations in the activity of SEF neurons (target onset: 270/822, 32.8%, **Fig. 4B**; initial choice: 306/822, 37.2%, **Fig. 4H**) predicted final choice. Among these neurons, a mean of 96.7 (SD=17.5) and 114 (SD=23.5) neurons predicted final choice in any given time bin when aligned to target onset and choice, respectively (**Fig. 4E, K**).

To estimate the magnitude of the influence of neuronal activity on final choice, we examined how the probability of choosing L in response to temptation (i.e., final L choice) changes for maximum and minimum activity levels of a neuron. **Fig. 4C, I** illustrate the model-predicted choice functions for what the final L choice would be when an SEF neuron’s activity was at its maximum (thick green line) or minimum (thick red line). For the neuron in **Fig. 4I**, higher firing rates in the relevant time bin (grey background) resulted in a final choice function that was shifted towards longer L delays, increasing the probability of finally choosing the L option. Neurons started predicting final choice early in the trial (target aligned: mean=-69.3ms, SD=6.97; choice aligned: mean=-51.5ms, SD=6.58), and could be either positively or negatively correlated with finally choosing L in response to temptation (**Fig. 4F, L**).

Altogether, 408/822 neurons (49.6%) predicted the final choice in at least one of the two alignments, the majority of which (239/408; 58.6%) had also been identified to encode self-control (**table S3**). 168/822 (20.4%) predicted final choice in both alignments.

## SEF activity and graded self-control

Our task design only allows us to distinguish between high and low self-control levels and we treated it accordingly as a categorical variable. However, self-control levels might vary continuously. Graded differences in self-control would not lead to behavioral differences in LL trials, since self-control above a threshold level should always lead to the rejection of temptation, independent of its strength. In contrast, in LS trials, the strength of self-control should be proportional to the length of time the monkey is able to withstand temptation before giving in to it (i.e., switch response time; switch RT) (**Fig. 5A**). There was considerable variation in the length of time the monkey waited before ultimately switching to the S option (**Fig. 2G**), which we used to search for graded differences in self-control levels.

**Fig. 5.**
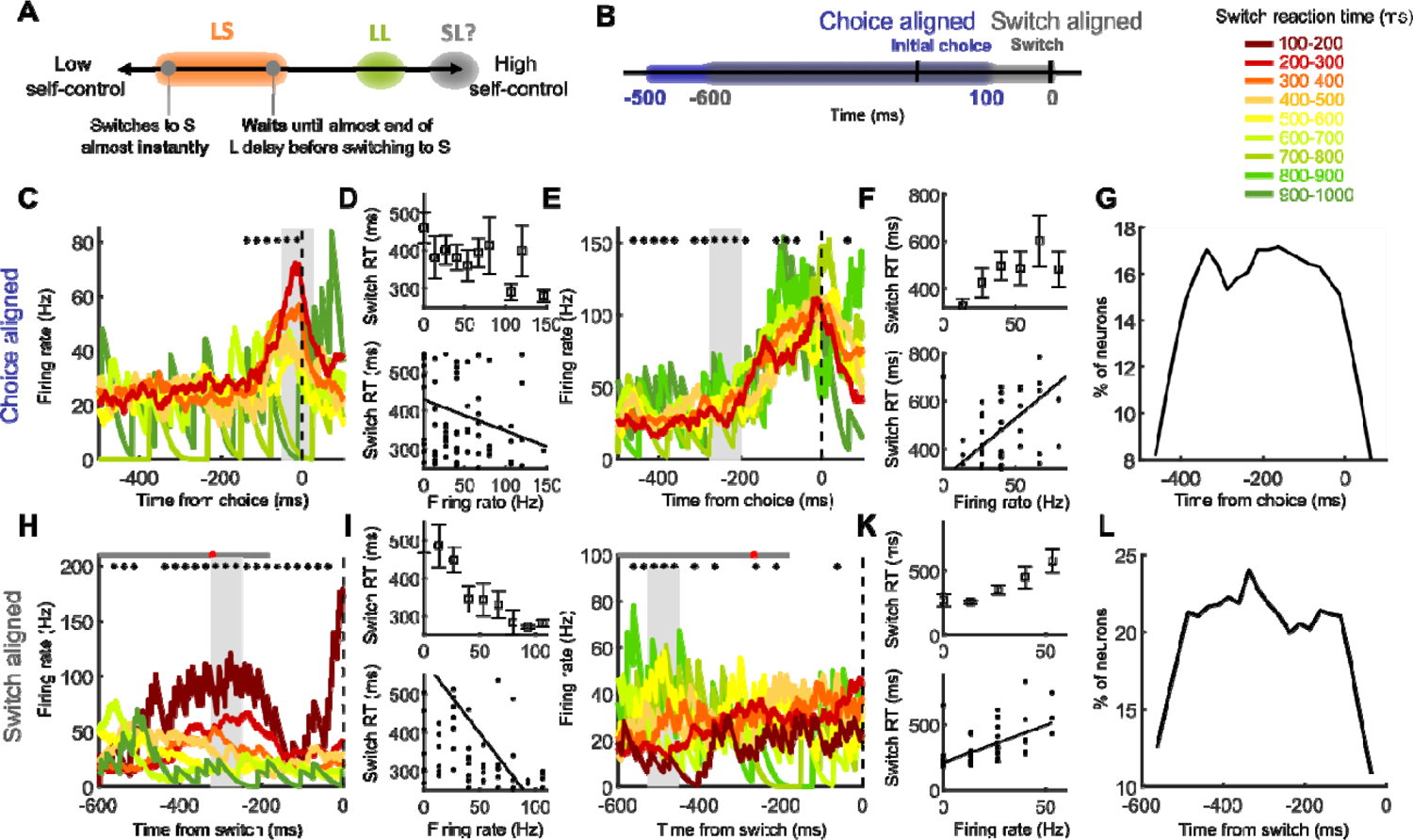
SEF neurons predict finer-grained differences in self-control. **(A)** Schematic illustration of a continuous gradient of self-control levels, and their potential behavioral consequences in the self-control task. **(B)** Analyzed time periods and their alignments. **(C)** Spik density functions for an example SEF neuron whose activity predicts and is negatively correlated with switch RT. Gray area indicates the time bin from which panel **D** was created. Stars indicate time bins in which the neuron’s activity predicted switch RT. Black bar indicates the range of switch RT in trials where this neuron was recorded. **(D)** Box plot (top) and scatter plot (bottom) of the relationship between switch RT and firing rate of the neuron in the 75-ms time bin. Boxes were visualized only if there was more than one data point per unique firing rate. **(E-F)** Same as in **C-D**, but for a neuron with a positive correlation with switch RT. **(G)** Proportion of neurons whose activity predicts switch RT, aligned to initial choice. **(H-L)** Same as in **C-G**, but aligned to switch. Black bars in **H**, **J** indicate the range of choice RT observed in trials the given neuron was active, and red dots indicate its median.

We tested whether SEF neuronal activity predicted switch RT. Only low self-control trials were used for this analysis. We examined neuronal activity in overlapping 75-ms time bins shifted by 25ms aligned to two events: (1) −500ms to 100ms around initial choice, and (2) −600ms to 0ms before switching (**Fig. 5B**).

A large proportion of SEF neurons (initial choice: 364 neurons, 44.3%; switch: 386 neurons, 47%) predicted switch RT (i.e., for 4 consecutive time bins). Across all time bins, an average of 140 (SD=24) and 162 (SD=28) neurons predicted switch RT when aligned to initial choice (**Fig. 5G**) and switch (**Fig. 5L**), respectively. Example neuronal activity correlated with switch RTs is shown aligned to initial choice (**Fig. 5C-F**) and switch (**Fig. 5H-K**). A substantial proportion of neurons that predicted switch RT was directionally tuned (choice: 281/364, 77.2%; switch: 292/386, 75.6%).

Overall, 509/822 neurons (61.9%) were identified to predict the timing of switch in at least one of the two alignments (**table S3**). The majority of these neurons (57%) also encoded self-control levels across all temptation trials. Thus, SEF activity not only differentiated between high and low self-control, but also reflected gradual differences in self-control.

## Discussion

To identify neural signals associated with self-control, we designed a novel task that reveals underlying self-control levels that are independent of cost-benefit evaluations. Previous studies identified a distributed network involved in self-control (*4*, *6*, *7*, *9*, *17*). The SEF, hypothesized to encode context-specific oculomotor action values (*18*, *19*), is interconnected with this network (*20*). Therefore, SEF is anatomically and functionally well-situated to encode and utilize self-control signals to causally influence behavior. Our work reveals the existence of self-control signals in the SEF. We focus here on activity preceding the initial choice. Because visual stimuli, choices, and motor activity are the same across high and low self-control trials until initial choice, it is unlikely that visual or motor differences caused the self-control signals.

Our findings argue directly against the hypothesis that self-control is merely a form of cost-benefit evaluation and does not require specific control circuits independent of value-based decision making (*10–12*, *21*). Under this hypothesis, activity differences between high (LL) and low (LS) self-control trials would be the result of fluctuations in the neural encoding of action value, which guides choice. However, a number of our findings argue against this interpretation.

First, in all LL and LS trials the monkeys initially preferred L over S, irrespective of possible preference variation across trials. This makes it difficult to explain subsequent switches as the result of neural activity differences at or before the time of initial choice, unless it represents correction of an error in the initial choice. Second, in the case of error correction, we would expect similar variability in action values for initial S choices, and likewise the same rate of SL switches. Instead, we observed dissimilar proportions of LS and SL switches (**Fig. 2B, C**). Third, the GLM analysis shows that SEF neurons carry neural signals for self-control, independent of signals related to value-based choice (e.g., chosen value, unchosen value) (**Fig. 3C**). If switches are the result of changes in value-based decisions, they should be accompanied by changes in subjective value, which would be reflected in the GLM through an interaction between self-control and value-related variables. However, we have clear evidence of SEF neurons encoding self-control without any such interactions **(table S5)**. Therefore, our results support the idea that self-control is independent of the valuation process, and can vary without altering the cost-benefit estimation process itself (*7*).

We concentrate here on self-control signals that emerge before the onset of temptation. This form of self-control represents proactive control (*22*, *23*). Proactive control maintains goal-relevant information before the occurrence of a cognitively demanding event, thereby anticipating and preventing interference to enhance performance. Such maintenance of goal-relevant information has been proposed to give rise to cognitive control by providing bias signals (*24*). Indeed, we found neurons that encoded the interaction between self-control and other variables that can be important for choice (e.g., spatial location of options, delay-discounted value of options) **(table S3)**. Moreover, 65.4% of SEF neurons predicting final choice also predicted initial choice. This suggests that self-control and value-based choice processes, while distinguishable, are not completely separated. Rather, it seems likely that self-control influences the value-based evaluation process that underlies the initial choice by biasing it towards the prudent option. This suggests that proactive self-control may already influence the likelihood of choosing the prudent option initially.

Thus, our results demonstrate that self-control is a process which is separable from value-based choice. SEF is involved both in explicit valuation judgments and self-control, which interact to influence initial and final choice, likely as one node in a larger network. These findings are in line with other results indicating a role of SEF in executive control (*18*, *19*).

## Materials and Methods

Five adult male rhesus macaque monkeys (*Macaca mulatta*) were trained to perform the self-control task (monkey A: 47 sessions; monkey I: 17 sessions; monkey G: 38 sessions; monkey K: 22 sessions, monkey H: 10 sessions). Overall, the five monkeys completed 108,568 trials over 134 behavioral sessions (no-temptation trials: 68,936; temptation trials: 39,632). Monkeys were surgically implanted with titanium head posts and a hexagonal titanium recording chamber. We recorded from a total of 822 Supplementary Eye Field neurons (monkey A: 208 neurons; monkey I: 102 neurons; monkey K: 366 neurons; monkey H: 146 neurons). All animal care and experimental procedures were approved by Johns Hopkins University Institutional Animal Care and Use Committee.

### Experimental Design

#### Self-control task

The search for neuronal self-control signals required a behavioral indicator that estimates the level of self-control on individual trials, independent from other factors that influence choice. Our self-control task provides such a behavioral indicator.

The self-control task (**Figure 1**) included two trial types: (1) no-temptation (46-90% of trials, mean across sessions = 63.5%) and (2) temptation (10-54% of trials, mean across sessions = 36.4%). We presented trials in blocks consisting of both trial types and all delay contingencies. Within a block, the order of appearance was randomized and a particular trial was never repeated, so that the monkeys could not anticipate the specific set of reward options before the targets were shown. Each target location was randomized and counterbalanced across trials to prevent the monkeys from preparing a movement toward a certain direction before target appearance.

The delay length of the S option (S delay) was fixed within each behavioral session (500 or 700ms), and the delay length of the L option (L delay) varied from trial to trial (500-10,000ms; equal to or longer than the S delay). The amount of reward for S and L options were fixed within the session for monkeys A, I, and G. For monkeys K and H, each session had two combinations of S and L rewards (e.g., [S reward, L reward]: [2,4] and [3,6]). For monkey H, the smaller reward combination did not produce a behavioral effect of temptation (**figure S3**). Therefore, for monkey H, trials that used the smaller reward combination within each session were not included for analysis.

Trials began with the appearance of a central fixation point. After the monkey maintained their gaze on the central fixation point (±2° window) for a variable delay (400-1,000ms), two choice targets appeared in locations opposite each other in one of four quadrants. Monkeys indicated their initial choice by making a saccade to one of the options and were required to fixate the chosen option cue (±1° cue, ±2° window) for the duration of the delay in order to be rewarded. If the S option was chosen, the intertrial interval was adjusted to match what the total duration of the trial would have been had the monkey chosen the L option. Therefore, to maximize total reward in a given session the monkeys should always choose the L option. However, choice behavior indicates that the probability of choosing the L option is dependent on the length of the L delay (**Figure 2A**), suggesting that the subjective valuation of the L option is discounted by the waiting time for L.

#### No-temptation trials

No-temptation trials were structured like the classic intertemporal choice task. A trial started with the monkey fixating a central fixation point. After a variable fixation period (400-1,200ms), two annulus-shaped stimuli were presented which served as saccade targets (**Figure 2B**). The orientation of the stimuli varied across trials. Simultaneously, the fixation point disappeared, which was a signal for the monkey to choose one of the targets. Each annulus had a colored section of variable length on it. The color of the section indicated the amount of reward and the length indicated the delivery delay associated with the target, respectively. The reward amount and the delay of the targets were adjusted so that one was always a small, short-delayed target (S option) and the other, a large, long-delayed one (L option). Once the monkey chose a target by fixating on it, the unchosen target disappeared, and the colored annulus section of the chosen target began to decrease with an angular velocity of 60°/sec (monkeys A, I), or in incremental ticks (monkeys G, K, H). On each trial, the initial angle of the colored section and the direction in which it decreases were randomized independently for each stimulus. If the monkey fixated the chosen target for the duration of the delay and its colored section was extinguished, fluid reward in the amount indicated by the color was delivered. If the monkey chose the S option, the intertrial interval was adjusted to increase by the difference between L delay and S delay. Thus, the total length of a trial was the same irrespective of the choice of the monkey. This adjustment ensured that there was no opportunity cost of choosing the L option.

The monkey was required to maintain fixation first at the fixation spot until it disappeared, and then at the chosen target until reward delivery (‘complete trials’). If the monkey broke fixation in either one of these two time intervals, the trial was aborted and no reward was delivered (‘incomplete error trials’). After the usual inter-trial-interval, a new trial started. For monkeys G, K, H, and later 13 sessions of monkey I (i.e., 83/134 sessions), an aborted trial was followed by a ‘repeat-trial’, which used the same S and L options as in the aborted trial. If the monkey aborted after making a choice, only the chosen target was presented, to create a forced-choice trial. If the monkey aborted before making a choice by breaking fixation from the fixation spot, both targets were presented. The location of the targets, however, was randomized so that the monkey could not prepare a saccade in advance. Such ‘repeat-trials’ were not included in the analysis.

#### Temptation trials

The initial sequence of events in temptation trials was identical to no-temptation trials. However, after the monkey made a saccade to one of the targets, the unchosen target did not disappear. Instead, both targets remained on the screen and the bands of both targets started to decrease simultaneously. For monkeys K and H, the unchosen option moved to the center of the screen. At this stage, the monkey could either maintain fixation at the initially chosen option (“stay trial”), or could change his choice by making a saccade to the initially unchosen option (“switch trial”). If the monkey switched, the initially chosen target disappeared. In trials where the monkey initially chose the L option, the band of the S option could extinguish before that of the L option. In this case, the S target changed to a filled circle indicating immediate reward availability. The intertrial interval was adjusted in the same way as in no-temptation trials depending on the final choice, ensuring that switching targets did not change the overall reward rate over all trials. Depending on which options were selected on the initial and final choice there were four different possible types of temptation trials: LL, LS, SS, and SL trials. On LL trials, the initial and final choices were the L option. On LS trials, the initial choice was the L option but the final choice was the S option. On SS trials, the initial and final choices were the S option. On SL trials, the initial choice was the S option but the final choice was the L option.

### Data Acquisition

#### Neurophysiological Recording Procedures

Spiking activity from single neurons were recorded extracellularly with single tungsten microelectrodes (impedance of 2-4 MΩs, Frederick Haer, Bowdoinham, ME) (monkeys A and I) and with multi-channel microelectrodes (16-channel S-Probes, Plexon Inc., Dallas, TX) (monkeys K and H). Electrodes were inserted through a guide tube positioned just above the surface of the dura mater and were lowered into the cortex under the control of a self-built microdrive system. The electrodes penetrated the cortex perpendicular to the surface of the cortex. Electrophysiological data were collected using a TDT system (Tucker-Davis technologies, Alachua, FL) at a sampling rate of 24.4 KHz. Action potentials were amplified, filtered, and discriminated conventionally with a time-amplitude window discriminator. Spikes were isolated online if the amplitude of the action potential was sufficiently above a background threshold to reliably trigger a time-amplitude window discriminator and the waveform of the action potential was invariant and sustained throughout the experimental recording. Spikes were then identified using a window discriminator (OpenEx, Tucker-Davis technologies, Alachua, FL) and the time stamps were collected at a sampling rate of 24.4 KHz. For monkeys K and H, action potentials from individual neurons were isolated and identified using principal component analysis (PCA) in the Plexon Offline Sorter (Plexon Inc., Dallas, TX).

#### Behavioral Recording Procedures

Custom MonkeyLogic software was used to control task events, stimuli, and reward, as well as to monitor and store behavioral events. During the experimental sessions, the monkey was seated in an electrically insulated enclosure with its head restrained, facing a video monitor. Eye positions were monitored with an infrared corneal reflection system, a 500 Hz video eye tracker (EyeLink 1000; SR Research, Ottawa, Canada) at a sampling rate of 1000 Hz. Reinforcement was delivered using a stepper motor-driven syringe pump fluid delivery system that is highly accurate across the entire range of fluid amounts used in the experiment. All analyses were performed using custom Matlab code, unless noted otherwise.

### Behavioral Analysis

#### Choice functions

The probability that the monkey chooses the S option was plotted as a function of the delay associated with the larger later reward (L delay). When the L delay was short, monkeys are more likely to choose the L option. As the delay of the L option increases, monkeys increasingly choose the S option. We used a logistic regression model to estimate the probability of choosing the S option, p(S), as a continuous function of the value of the L delay (i.e., the choice function):

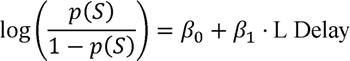

For no-temptation trials, the choice function was estimated for only the initial choice because once an option was selected and the monkey fixated on its choice for the required amount of time, the trial ended. For temptation trials, the choice function was estimated for both the initial and the final choice. The L delay at which choosing either the L or S option is equal [p(S) = 0.5] is known as the indifference point (*2*). By definition, at this point, the subjective value of the two options must be equal. We called this the indifference delay D_indiff_ (i.e., the L delay at which the monkey is indifferent between the S and L options). We used D_indiff_ to estimate the temporal discounting parameter and the subjective value of each option. Because monkey K had two distinct combinations of S and L reward amounts interleaved within each session, the indifference delay was computed separately for each reward combination.

#### Estimation of Subjective Value (SV)

SV of options was calculated by delay-discounting the final reward amount, using the following hyperbolic discounting formula (*22*):

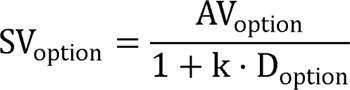

SV_option_ is the calculated SV of the option, AV_option_ is the actual reward amount the monkey is to receive, k is the delay discounting parameter which reflects how much the animal discounts later rewards, and D_option_ is the length of delay of that option. The discounting parameter, k, was estimated by fitting a logistic regression model using maximum likelihood estimation. Note that the term value, unless noted otherwise, refers to the estimated delay-discounted subjective value (SV), and not the absolute reward amount.

### Neurophysiological Analyses

We used generalized linear models (GLM) (for encoding and decoding models) and linear models (for graded self-control models) for neural analyses. Only completed (i.e., trials where the monkey did not break fixation from the chosen option until the end of the delay and thus received a reward) no-temptation and temptation trials were used in neural analyses.

#### Encoding models

Encoding models were GLMs designed to predict SEF neuronal activity (*Spike count*) from a series of task-related variables in three different time periods: (1) *central fixation*: 0 to 400ms around fixation onset (i.e., the time the monkey fixated on the central fixation point), (2) *target onset*: −50 to 300ms around target onset (i.e., when choice targets appeared), (3) *before choice*: −300 to 50ms around initial choice.

To quantitatively characterize how each SEF neuron’s activity is modulated by task-related variables in different time periods throughout the task, we performed Poisson-GLM analyses (*23*). Because the Poisson distribution is used to model the number of times an event occurs in a given interval of time, it can also be used to characterize the firing activity of a neuron. Under the assumption that neuronal firing (i.e., the number of spikes) follows a Poisson distribution, the activity of a neuron in a given time period can be represented as the exponential of the linear combination of predictor variables.

Within each time period, the full model including all variables and each of its nested models, including the null model, were fit in 75-ms moving windows, shifted by 25ms. The model with the lowest Akaike Information Criterion (AIC) was selected as the neuron’s best fitting model in the given time bin. The predictors included in the best fitting model were regarded to be represented, or encoded, by the neuron in that time bin.

For all time periods, the full model included a constant term, 5 choice-related variables, the variable *Self-control*, and select 2-way interactions (**table S2**):

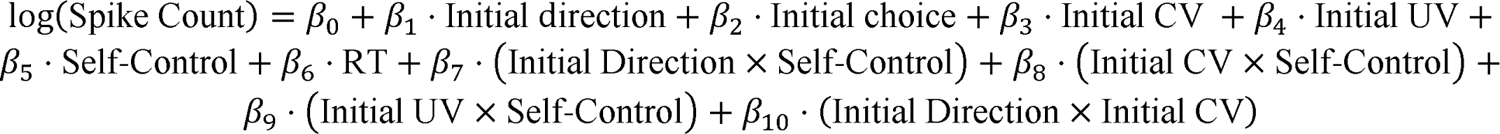

*Initial direction* was a categorical variable that indicated the location of the chosen target to which the saccade was directed. There were four possible target locations, one in each quadrant. *Initial choice* was another categorical variable which indicated the identity of the chosen option, i.e., S or L options. There were two value-related variables: *Initial chosen value (Initial CV)* was defined as the subjective delay-discounted value of the chosen option in the given trial, and *Initial unchosen value* (*Initial UV*) was the subjective value of the unchosen option in the given trial. Because we found a small but significant difference in choice reaction time (choice RT) for high (LL) and low (LS) self-control trials (mean RT difference LS-LL; monkey A: 22.5ms, I: 23.7ms, G: 0.5ms, K: 4.79ms, H:-6.69ms; Kolmogorov-Smirnov test, k=0.0787, p=5.68e-59) (fig. S5), we included *RT* as a potential variable to dissociate the effects of self-control and motor preparation on explaining neuronal activity.

*Self-control* was a categorical variable which distinguished amongst trials with high self-control, low self-control, and other trials. High self-control trials were defined as temptation trials in which the monkey initially chooses the L option and stays with it (LL trials) and low self-control trials were defined as temptation trials in which the monkey initially chooses the L option but switches to the S option (LS trials). All other correct trials, that is: (1) no-temptation trials, (2) temptation trials in which the monkey chose and stayed with the S option (SS trials), and (3) temptation trials in which the monkey chose the S option but switched to the L option (SL trials) were considered as trials with an undetermined self-control level. In no-temptation and LS trials, it was impossible to distinguish high and low self-control levels. In SS trials, self-control was not required to reach the end of the trial and receive the reward.

The response variable, *Spike Count*, was defined as the number of action potentials fired by the neuron within the given time bin. For visualization in figures, spike count was later converted to firing rate (*FR*; spikes/sec).

#### Coefficient of Partial Determinantion (CPD)

To quantify the unique contribution of each predictor on SEF neuronal activity over time, the mean CPD across neurons was computed for each predictor in each time bin. CPD reflects how much variance in a neuron’s activity is uniquely explained by each variable, relative to that neuron’s best fitting model at a given time bin. Using the best-fitting model of a neuron as its “full model”, we calculated CPD for a given predictor as:

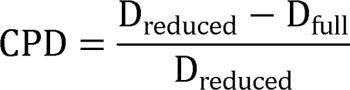

D_full_ and D_reduced_ are the deviance values of the best-fitting model and the reduced model that excludes the given predictor from the best-fitting model, respectively. For visualization, outliers were excluded using MATLAB’s *rmoutliers* function before averaging.

#### Decoding models

Decoding models were GLMs designed to predict choice from SEF neuronal activity (*spike count*) in two time periods: (1) −200 to 300ms around target onset, and (2) −200 to 300ms around initial choice. Within each time period, the full model including all variables and each of its nested models were fit in moving windows of 75-ms time bins, shifted by 25ms. For both time periods, the predictors other than *spike count* were *L delay* and *initial direction*, where *L delay* refers to the L option’s delay length and *initial direction* refers to the spatial location of the initially chosen option. This was done both for predicting initial choice and final choice. Since *L delay* is closely correlated with the subjective value of the L option and is known behaviorally to influence choice, the activity of neurons that represents the decision process would not be expected to explain variance in final choice beyond the influence of *L delay*. In contrast, a neuron whose activity explains a significant proportion of the variance in final choice beyond the influence of *L delay* should result in time bins where the model includes the predictor *spike count*. For such a neuron, increasing firing rate should be correlated with either a rightward shift or leftward shift in the choice functions. The inclusion of saccade direction as a predictor allowed us to determine if the influence of neuronal activity was spatially tuned.

For predicting initial choice, *Spike Count*, *L Delay*, and *Initial Direction* from both no-temptation and temptation trials (L, S, LL, LS, SL, SS trials) were used as predictors of the model. For predicting final choice, *Spike Count*, *L Delay*, and *Initial Direction* from only the temptation trials (LL, LS, SL, SS trials) were used as predictors of the model. The full model includes all of the predictors, a constant term, and all two-way and three-way interactions of the predictors:

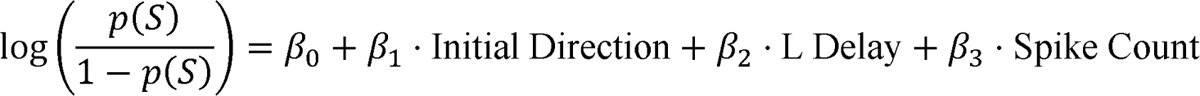

For each time bin for a given neuron, the full model and its nested models were fit to the choice data. The model with the lowest AIC was taken as the neuron’s best fitting model. A given neuron was interpreted to influence choice over and above the behavioral variables only if the predictor *spike count* was included in the best fitting model.

#### Graded self-control model

Graded self-control models were linear regression models designed to predict the latency for switching (*Switch RT*) from *Spike Count* of the SEF neuron, controlling for other task-related variables that can influence switch latency in the given trial. We examined two time periods: (1) −500 to 100ms aligned to initial choice, and (2) −600 to 0ms aligned to time of switch. Within each time period, the full model including all variables and each of its nested models were fit in moving windows of 75-ms time bins, shifted by 25ms.

Switch response time (*Switch RT*), defined as the time between initial choice and time of switch, was the response variable. Because Switch RT exhibited a right-skewed distribution (**Figure 2G**), we applied a log-transformation to better meet linear model assumptions. Thus, all models were fit using the log-transformed Switch RT as the dependent variable. Predictors other than *Spike Count* (number of spikes by a given neuron in the given time bin) included *CV-UV* (difference between the subjective values of the chosen and unchosen option) and *Initial Direction* (location of the chosen option). This aimed to capture neurons whose activity around initial choice significantly predicted the time the monkey waits after realizing it is a temptation trial (i.e., the unchosen option does not disappear and remains available) before ultimately switching. Only temptation trials where the monkey initially chose the L option (i.e., LS trials) were included for analysis.

The full model includes the above predictor variables, a constant term, and all two-way and three-way interactions, producing the following formula:

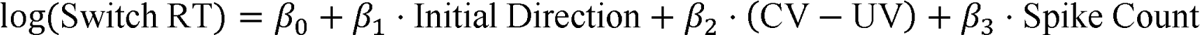

The *CV-UV* variable can influence the decision of whether and when to switch. Therefore, a neuron whose activity around initial choice significantly influences variance in switch latency beyond the influence of *CV-UV*, should have a best fitting model (defined as the model with the lowest AIC) that includes *Spike Count* as a predictor.

## Acknowledgments

The authors would like to thank Brance Amussen, Arless Carpenter, Ofelia Garalde, Justin Killebrew, Bill Nash, and Bill Quinlan for technical help with surgery, the experimental setup, and the computational infrastructure.

## Author contributions

Conceptualization: JH, KL, VS Methodology: EE, JH, KL, VS Investigation: EE, JE, JH, KL Visualization: KL

Funding acquisition: VS Project administration: VS Supervision: VS

Writing – original draft: KL, VS Writing – review & editing: KL, VS

## Supplementary Materials

**fig. S1.**
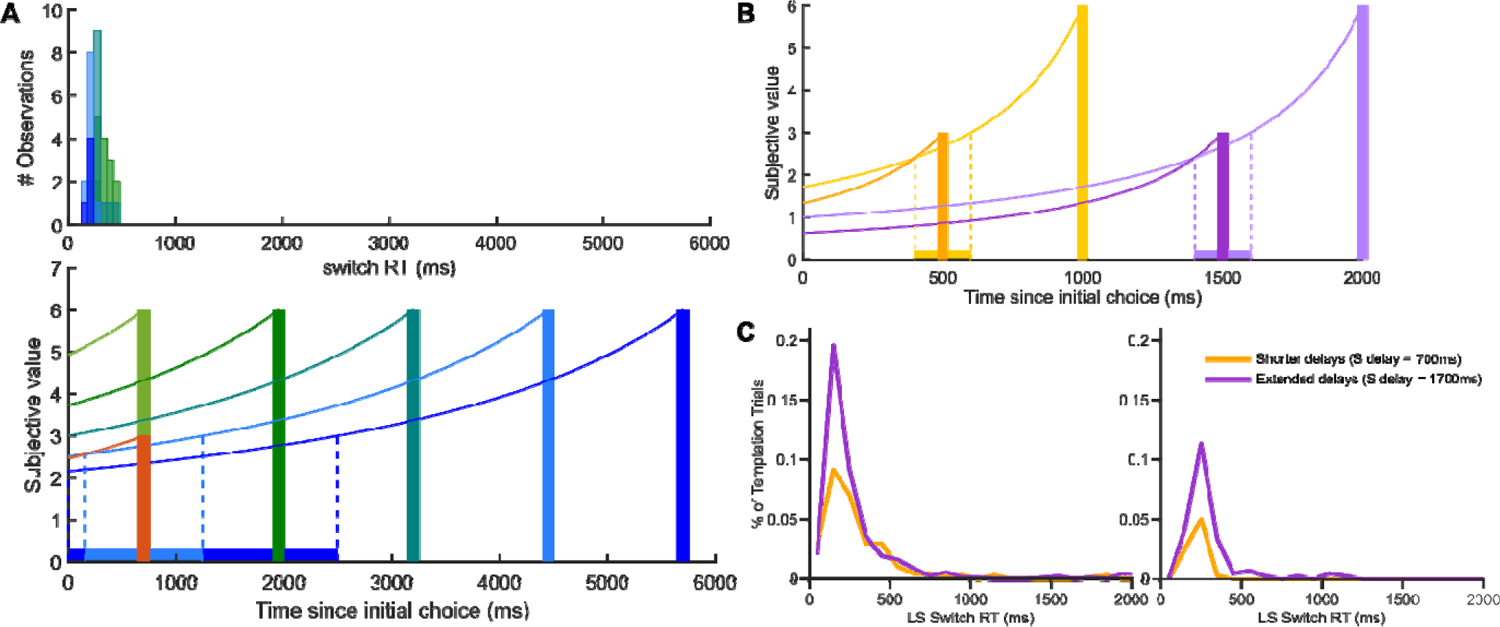
Empirical evidence that argues against the hypothesis that switches are byproducts of temporal discounting. **(A)** (Top) Distribution of the timing of switching from the L to the S option (switch RT) after initial choice, for each unique L delay in an example behavioral session. (Bottom) Estimated subjective value of S and L rewards in the example behavioral session. The momentary subjective value reflects hyperbolic discounting based on the remaining waiting time until reward delivery. The discounting parameter, k, was estimated from choice behavior in the session. The observed switch RT distribution is not incompatible with the hypothesis that switches result from temporal discounting, especially if one allows for random fluctuation of the neuronal representations. However, some inconsistencies are also present. If switches are driven primarily by the transient reversal of delay-discounted S and L values, they should occur within the range of time where S value exceeds L value (colored regions within dotted lines), with a peak at where the value difference is maximal. Instead, switches are observed at timepoints where the S value does not exceed L value, and the observed switches occur earlier than is predicted by the hypothesis. Note that for the 3 shortest L delays (light green, dark green, teal), the hypothesis would predict that no switches occur. However, a considerable number of switches are observed. (**B)** Hypothetical hyperbolic discount functions of subjective value for a given set of S (orange) and L (yellow) options and the predicted range of switch RT (colored yellow region within dotted lines). The dark (S) and light (L) purple lines indicate resulting hyperbolic discount functions of the same options after adding one second to both S and L delays. The predicted range of switch RT (colored light purple region within dotted lines) for these altered discount functions indicates that adding a time interval of equal length to both S and L delays should (1) push the switch RT distribution to later in time, but (2) should not change the frequency of switches. **(C)** Observed switch RT distributions (pooled over all sessions) for monkeys K (left, 9 sessions) and H (right, 4 sessions), before (yellow) and after (purple) adding one second to both S and L delays. We (1) did not observe a shift in the switch RT distribution, and we (2) observed a stark increase in switches for all L delays. This suggests that a dynamic readjustment of the values of L and S in the waiting period following initial choice is not the primary driving force of switches.

**fig. S2.**
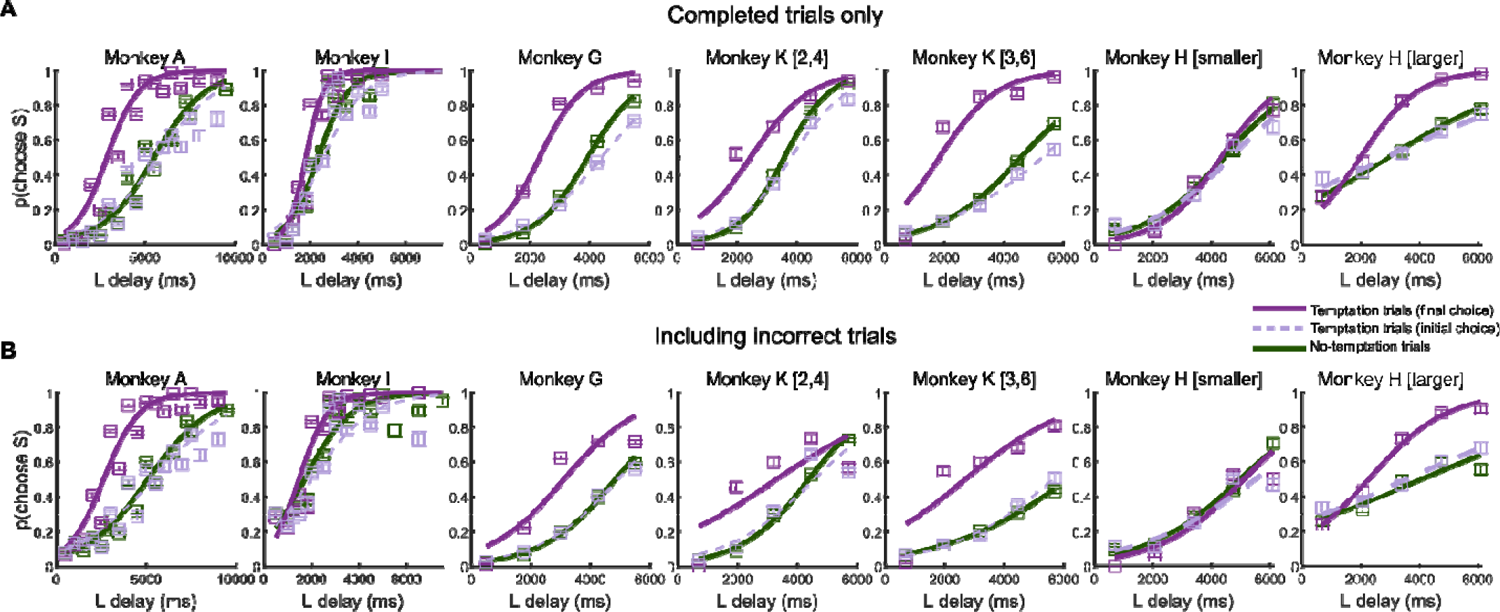
Effect of aborted trials on choice functions. Monkey K had two combinations of S and L reward amounts interleaved in each session. In the main figure, for monkey K, only the larger reward combination (S reward = 3, L reward = 6) was visualized (but all trials were included in analyses). Here, we show the choice functions for both reward combinations. Monkey H also had two combinations, but because the smaller reward combination did not yield a behavioral effect, all trials with the smaller reward combination were removed from all behavioral and neural analyses. **(A)** Choice functions estimated by using completed trials only (i.e., the monkey did not break fixation and received a reward). The initial choice function of no-temptation trials has a steeper curve than that of temptation trials, a difference more pronounced at the longer L delays. This apparent systematic difference in delay attitude is likely the result of only using completed trials in analyses. By design, when the monkey makes an L choice in no-temptation trials, the monkey must remain fixated until the end of the delay in order to be rewarded. If the monkey breaks fixation, the trial is aborted and is not included in analyses. However, in temptation trials, after the monkey makes an L choice, the monkey can break fixation to settle for the S option, such that the trial is completed and becomes included in analyses. Thus, a higher proportion of initial L-choice trials were included in analysis for temptation trials than for no-temptation trials (mean initial L-choice proportion for no-temptation trials: 62.5%, temptation trials: 64.5%; paired t-test, t=-3.59, p=4.58e-04). This also explains why the difference in initial choice functions between no-temptation and temptation trials is more pronounced for longer L delays. Such a difference in initial L-choice trials for temptation and no-temptation trials were not observed when incomplete trials were included (mean initial L-choice proportion for no-temptation trials: 54.6%, temptation trials: 55%; paired t-test, t=-0.57, p=0.57). **(B)** Choice functions estimated by including incomplete trials (i.e., including aborted trials where the monkey broke fixation before the end of delay and was not rewarded). Choice functions that include all trials, regardless of whether they were completed, exhibit no such significant difference in initial choice between no-temptation and temptation trials (paired t-test, t=0.7335, p=0.4644).

**fig. S3.**
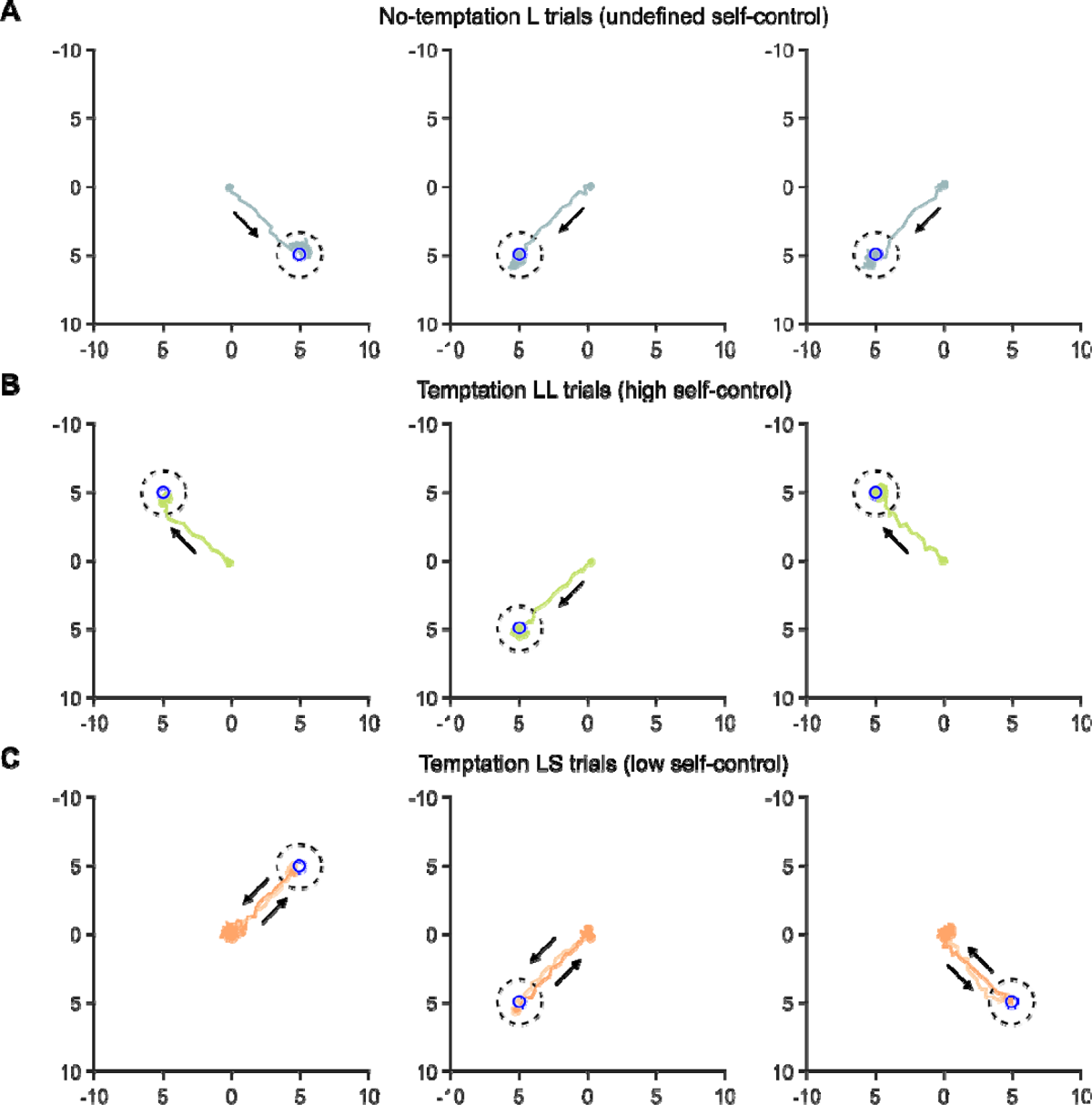
Example eye traces for L, LL, and LS trials. Blue circle indicates the location of the L option. Black dotted circle indicates the fixation window within which the monkey was considered to be fixating on the target. Arrows indicate the direction of saccades. Axes units are in degrees of visual angle. The central fixation point was presented at (0,0). **(A)** Eye traces in three example no-temptation L trials. **(B)** Eye traces in three example temptation LL trials. **(C)** Eye traces in three example temptation LS trials (i.e., switch trials). Because these trials are from monkey K, the location of the unchosen S option after the initial choice was moved to the center of the screen (0,0).

**fig. S4.**
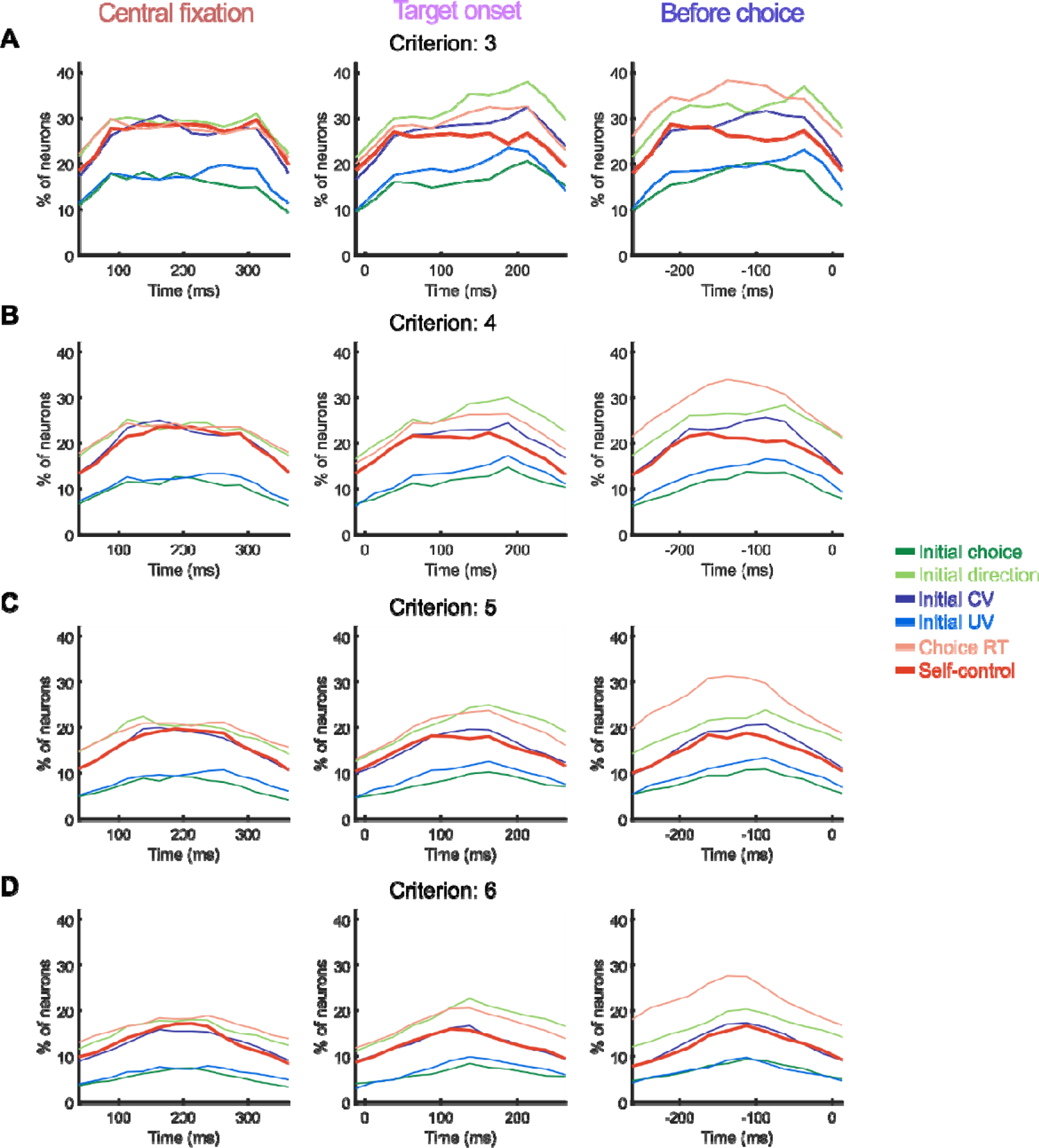
Effect of criterion condition for counting a neuron as encoding a variable. In the main text, the criterion for including a given variable in a neurons best-fitting model was set to be 4 consecutive time bins. Here we show the effects of using more or less strict conditions. (**A**) The criterion is set more lenient to 3 consecutive time bins. **(B)** The criterion is set to 4 consecutive time bins, as used for the main text (see Figure 3C). **(C-D)** The criterion is set more strictly to 5, and 6 consecutive time bins, respectively. The overall pattern is shown to be qualitatively unchanged, regardless of the criterion used.

**fig. S5.**
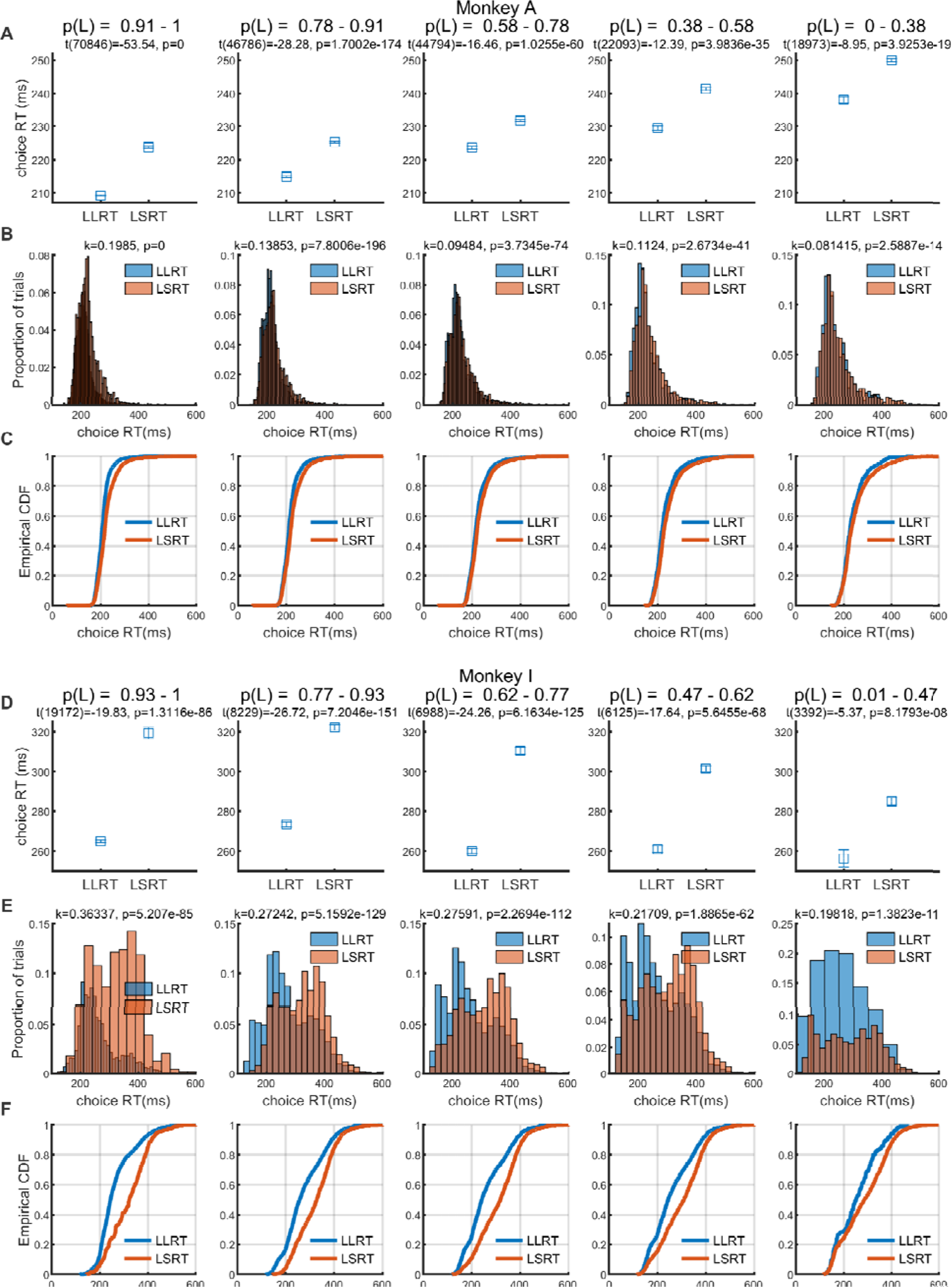
Systematic difference in choice reaction time (choice RT) between LL and LS trials. Monkeys A, I, G, and K on average initially chose the L option more quickly (and monkey H more slowly) when they would resist temptation (LL) than when they would give in to temptation (LS) (mean RT difference_LS-LL_; monkey A: 22.5ms, I: 23.7ms, G: 0.5ms, K: 4.79ms, H:-6.69ms; Kolmogorov-Smirnov test, k=0.0787, p=5.68e-59). This difference persisted even after controlling for the subjective value of the L option, suggesting that whether the monkey would overcome temptation is at least partially associated with the speed at which the monkey chooses the L option. This difference in choice RT for LL and LS trials can be driven by different levels of saccade vigor, which could reflect a combination of self-control, action value, or lower-level saccade mechanics such as random fluctuations in the excitability of saccade planning circuits. Here, we illustrate the systematic difference in choice RT between LL and LS trials for the two monkeys that exhibited greatest difference. **(A)** Mean choice RT for LL (high self-control) trials and LS (low self-control) trials in monkey A, controlling for subjective value, reflected by the probability of choosing L (p(L)). Because subjective valuation for a given L delay could differ from session to session, we calculated the probability of choosing the L option for each L delay separately for each session. Trials were grouped into five percentiles by the computed p(L) to plot the five panels from high L value (leftmost) to low L value (rightmost). Titles report results from the two-sample t-test. Note that choice RT for LL trials (LLRT) is significantly lower than that for LS trials (LSRT) in all five groups. **(B)** Distribution of choice RT for LL and LS trials, plotted separately for each group. Titles report results from the Kolmogorov-Smirnov goodness-of-fit test. **(C)** Empirical cumulative density functions of choice RT in LL and LS trials, plotted separately for each group. **(D-F)** Same as in **A-C**, but for monkey I.

**fig. S6.**
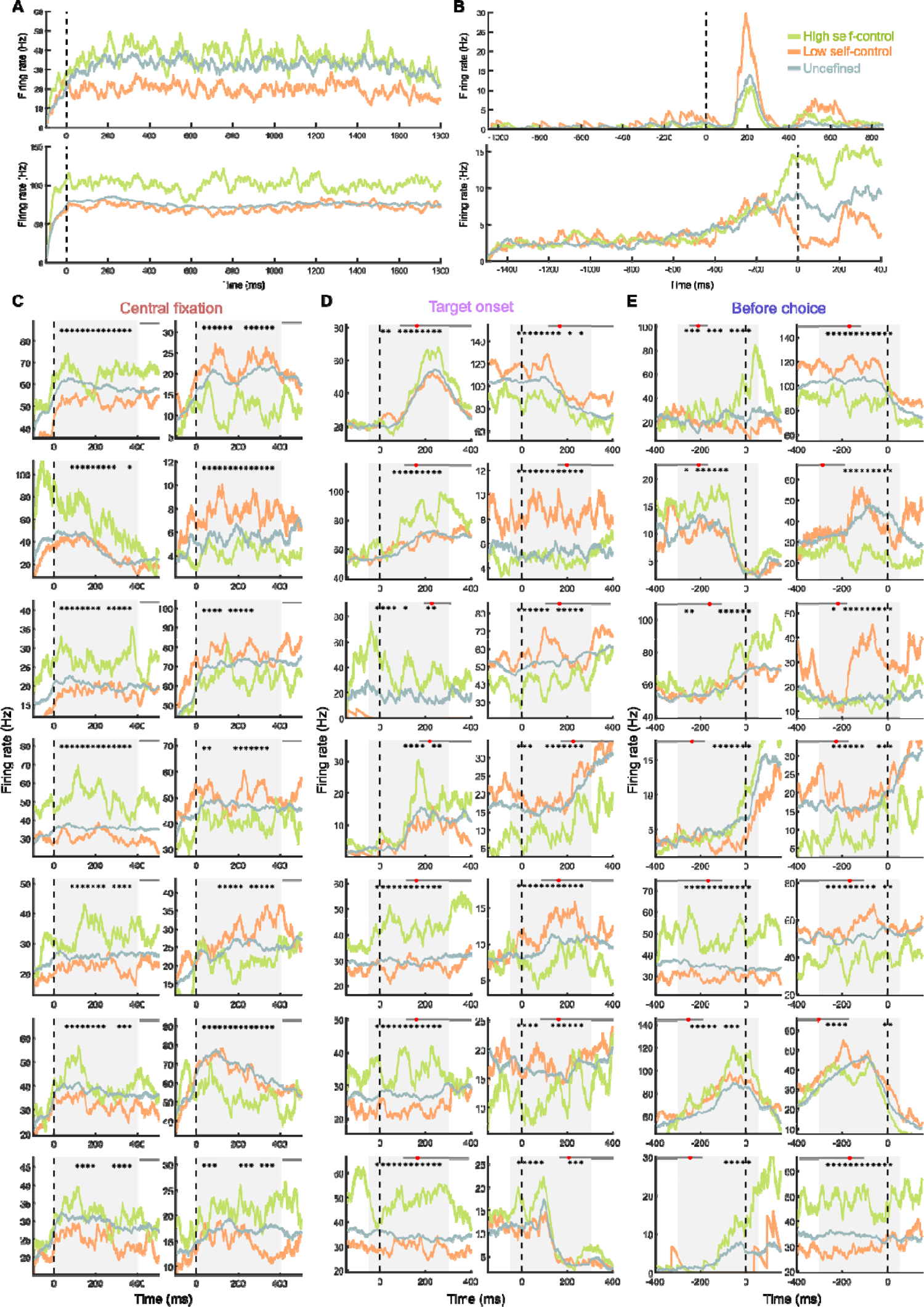
Additional example neurons encoding self-control. **(A)** Two example neurons that exhibit sustained activity differences for high and low self-control, both aligned to fixation onset. **(B)** Two example neurons that exhibit transient activity differences for high and low self-control, (top) aligned to target onset, (bottom) aligned to initial choices. **(C-E)** Additional example neurons that encode self-control during central fixation **(C)**, after target onset **(D)**, and before choice **(E)**.

**fig. S7.**
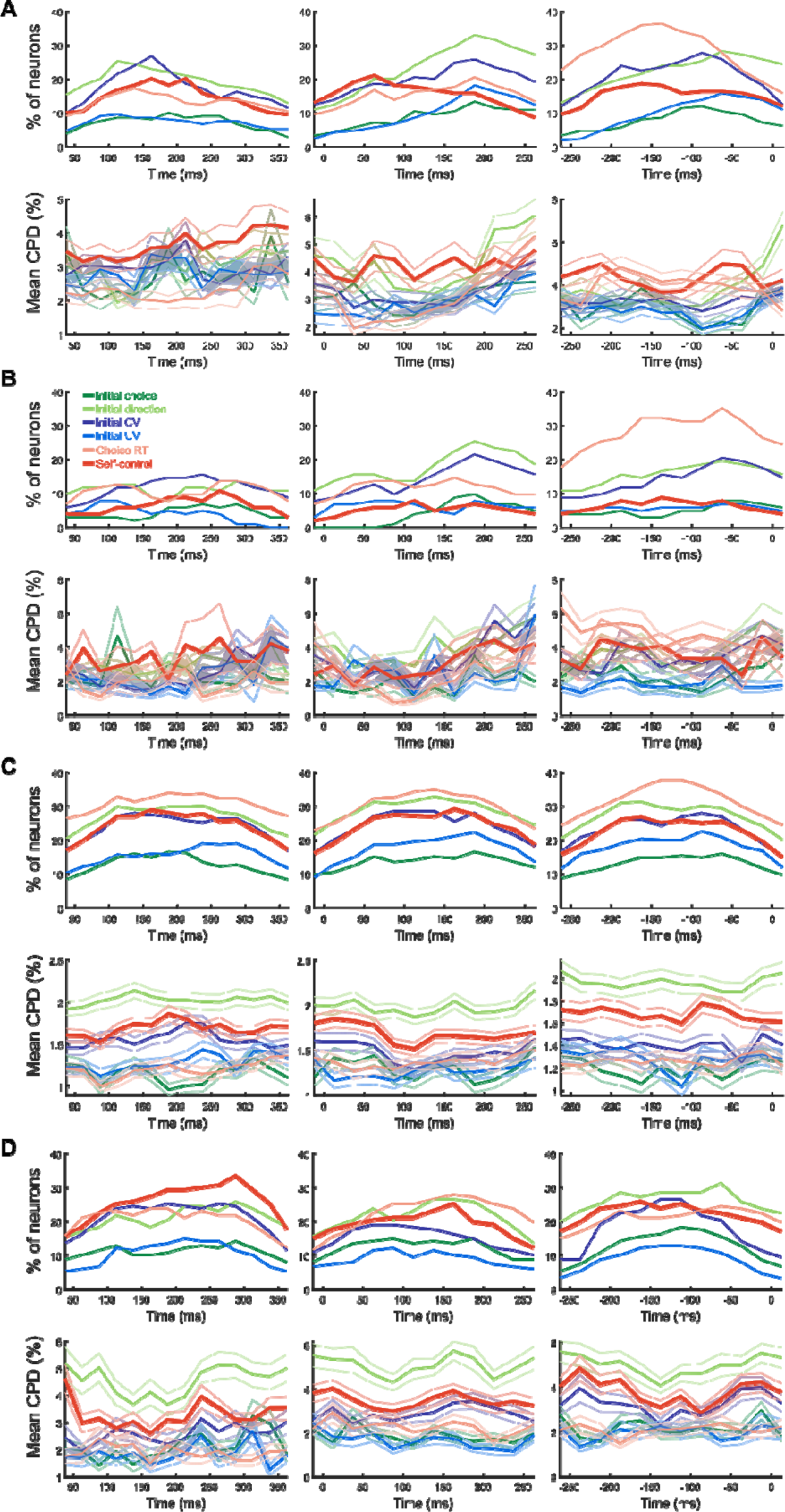
Neurons encoding self-control and mean CPD separately by monkey. **(A)** Proportion of neurons from monkey A, encoding each variable in a given time bin. (bottom) Mean coefficient of partial determination (CPD) for each variable in each time bin. The shaded areas represent SEM. **(B-D)** Same as in **(A)**, but for neurons from monkeys I, K, and H, respectively.

**fig. S8.**
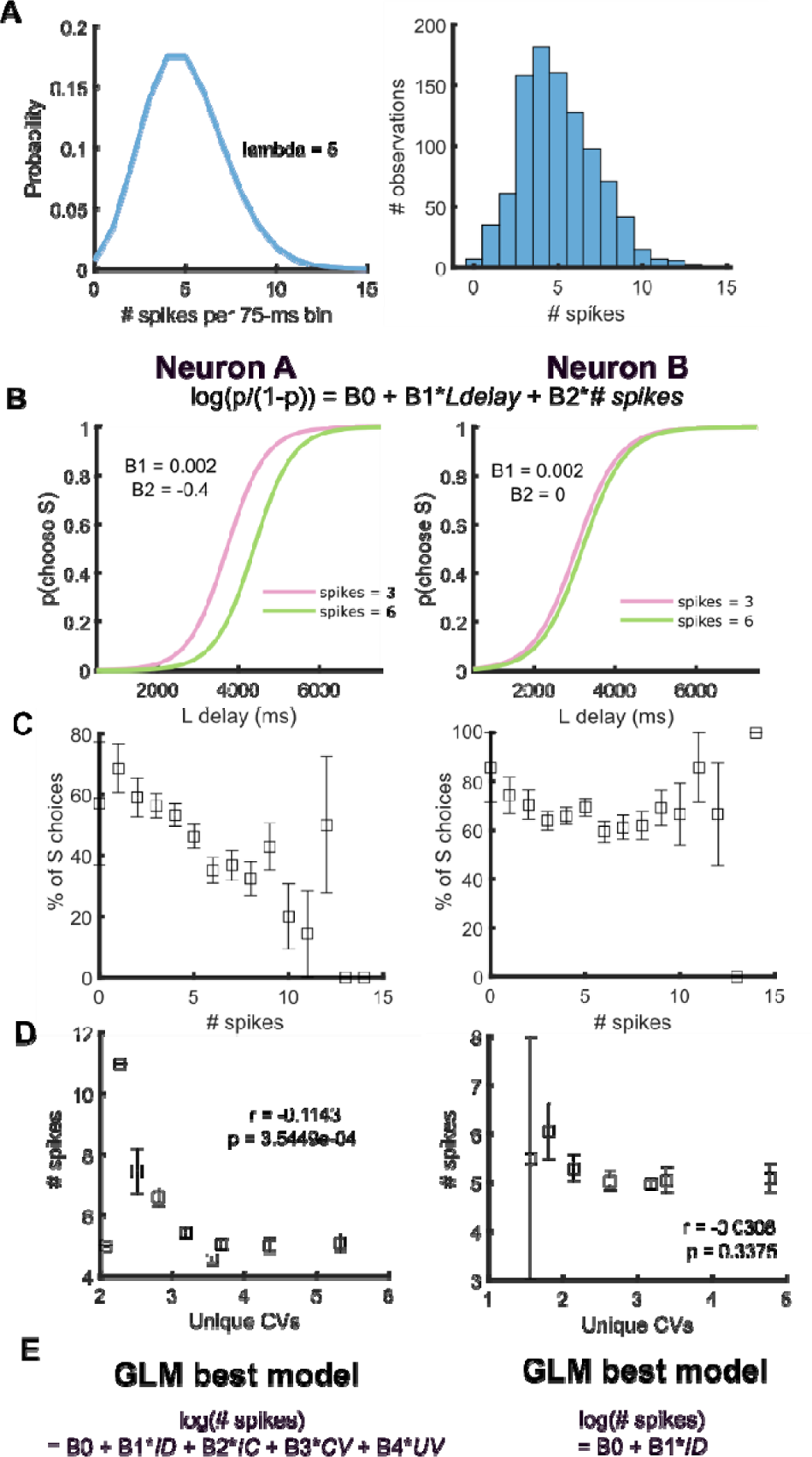
Hypothetical mechanism whereby a proactive self-control signal can lead to a spurious correlation with choice-related variables during the fixation period. We found that SEF neurons encode initial choice (S or L option) during the fixation period at the beginning of the trial, even before the targets are presented. While at first perplexing, the relationship becomes more understandable, if one considers that the task structure ensures that in every trial a S and a L option is presented. It is therefore possible that the monkey already has a bias for selecting one of these options or the other at the very beginning of the trial. Such a bias would not necessarily completely determine the later choice, but it could shift the fraction of L or S choices for a particular pair of options. The bias signal for the initial choice can be interpreted as a form of self-control. In the beginning of a trial, the monkey (by virtue of the task structure) knows S and L options will be presented and can decide to commit to a particular option prior to target presentation, and these SEF neurons act to bias the monkey towards a particular option reflecting that pre-target decision. In the main paper we did not include the initial choice of L as reflecting self-control because it could be confounded with cost-benefit analysis processes. However, this does not preclude the possibility that self-control is also involved in this initial choice of L. From this perspective, the presence of these neurons, before target presentation, which appear to be correlated with choice, seem to support the hypothesis that self-control also plays a part in the initial choice of the prudent option. Of course, other factors unrelated to self-control could also contribute to a choice bias. What is harder to understand is that the neuronal activity also reflects initial choice-related signals during the fixation period, such as chosen value (CV), or unchosen value (UV), since the information on which these choice variables depend is only revealed at target onset. However, if a neuron’s activity is correlated with upcoming choice, its activity can also be correlated with chosen value (CV). This is true regardless of which specific combinations of options are presented later in a specific trial. As long as neural activity is correlated with initial choice (i.e., biases toward a particular choice option), its activity can be shown to be also correlated with CV, because CV is correlated with initial choice. In this figure we illustrate, how a hypothetical neuron encoding choice bias could also show spurious correlations with other choice-related variables. **(A)** A hypothetical neuron whose trial-to-trial activity (in number of spikes) in a given 75-ms time bin is randomly sampled from a Poisson distribution. (left) The Poisson probability density function, with lambda = 5. (right) Histogram of sampled # spikes from the Poisson distribution, used to simulate the spiking activity of two hypothetical neurons. **(B-E)** Functionally, the choice bias can be modeled using a GLM that explains choice as a function of the L delay and spiking rate. The relative strength of the neuron’ s biasing effect is expressed as the relative strength of its influence on predicting initial choice (i.e., magnitude of the coefficient for neural activity; B2 in panel B). In the following, we illustrate the relationships of two hypothetical neurons A and B with choice behavior. The spiking activity of both neurons is derived from the same Poisson probability density function shown in A. Hypothetical neuron A’s activity is correlated with upcoming initial choice (B2 = −0.4; higher activity increases the probability of a L choice). In contrast, hypothetical neuron B shows the same spiking patterns, but the neuronal activity is not correlated with initial choice (B2 = 0). Only neuron A’s activity is expected to be correlated with the choice pattern. Neuron B serves as a control. We used the Poisson distribution and the GLM models to simulate neuronal activity and choices for each neuron. The following plot show the results of analyzing these simulated data sets. (**B**) The red and green choice functions indicate the probability of a S choice as a function of the later revealed L delay, when the hypothetical neuron A (left) or neuron B (right) producing 3 and 6 spikes in a given time bin in the fixation period. **(C)** Box plots showing the relationship between neural activity of neurons A and B in the fixation period and the overall proportion of S choices. AS expected, neuron A’s, but not B’s, activity is negatively correlated with S choice. **(D)** Box plots showing the relationship between neural activity of neurons A and B in the fixation period and initial chosen value (CV). Note that neuron A’s activity is significantly correlated with CV. r denotes the Pearson correlation coefficient. **(E)** The best-fitting models for hypothetical neurons A and B. Neuron A, whose activity was designed to be correlated with choice is found to encode initial choice (IC), but also CV and UV. Because neural activity is parametrically related to initial choice (i.e., stronger activity leads to a stronger tendency towards choosing a particular option), there can be cases where neural activity is better explained by a continuous variable (e.g., CV) than a binary variable (e.g., initial choice). Thus, neurons that were found to “encode CV” during fixation where monkeys could not have known the delay-discounted value of choice targets, could be neurons whose activity is related to the upcoming choice.

---

**table S1.** Variables and interactions considered for GLM. Variables and select two-way interactions included for encoding models.

| Predictors |
| --- |
| Self-control |
| Initial direction |
| Initial choice |
| Initial chosen value (Initial CV) |
| Initial unchosen value (Initial UV) |
| Choice RT |
| Initial direction * self-control |
| Initial CV * self-control |
| Initial UV * self-control |
| Initial direction * initial CV |

**table S2.** Number of neurons positively or negatively correlated with self-control in different time periods. A neuron is counted as purely positive or negative if it encodes self-control and its activity was positively or negatively correlated in all time bins that included self-control in the neuron’s best fitting model. Some neurons showed a positive correlation in some time bins and a negative correlation in others. Such neurons are counted as mostly positive (or negative) if they encode self-control and their activity was positively (or negatively) correlated with self-control in the majority of time bins that included self-control in the neuron’s best fitting model. Note that purely positive (or negative) neurons and mostly positive (or negative) neurons are mutually exclusive categories. The proportion of negatively and positively correlated neurons was similar in all time periods (one proportion z test: central fixation: z=0.0576, p=0.954; after target: z=-1.27, p=0.204; before choice: z=-0.816 p=0.415).

| Correlation with self-control |  | Number of neurons (n=822) |  |  |
| --- | --- | --- | --- | --- |
|  |  | Fixation | Target | Choice |
| Positively correlated | Purely positive | 66 | 55 | 58 |
|  | Mostly positive | 85 | 61 | 64 |
|  | Total | 151 | 116 | 122 |
| Negatively correlated | Purely negative | 52 | 50 | 56 |
|  | Mostly negative | 98 | 97 | 86 |
|  | Total | 150 | 147 | 142 |
| **Total** |  | **301** | **263** | **264** |

**table S3.** Number of self-control related neurons identified by each analysis, separately by monkey. For each row, the number indicates the number of neurons identified by each analysis, separately by monkey. The total number of neurons recorded from each monkey is indicated in brackets.

| Analysis | Number of neurons (n=822) |  |  |  |  |
| --- | --- | --- | --- | --- | --- |
|  | Monkey A<br>(n=208) | Monkey I<br>(n=102) | Monkey K<br>(n=366) | Monkey H<br>(n=146) | Sum<br>(n=822) |
| Encoding of self-control | 100 | 27 | 231 | 85 | 443 |
| Prediction of final choice | 112 | 45 | 194 | 57 | 408 |
| Prediction of switch RT | 128 | 48 | 252 | 81 | 509 |

**table S4.** Encoding of other variables and interactions. A neuron is counted as encoding a variable or interaction if the variable or interaction included in the neuron’s best fitting model for 4 consecutive time bins (i.e., the signal is present for 150ms). The count for variables includes neurons that encode interactions (i.e., a neuron that encodes the initial direction * self-control interaction is counted as encoding initial direction, self-control, and initial direction * self-control).

| Variable or interaction | Number of neurons (n=822) |  |  |
| --- | --- | --- | --- |
|  | Fixation | Target | Choice |
| Initial direction | 316 | 341 | 328 |
| Initial choice | 177 | 176 | 163 |
| Initial chosen value | 321 | 301 | 304 |
| Initial unchosen value | 196 | 205 | 209 |
| Choice RT | 302 | 291 | 348 |
| Initial direction * self-control | 217 | 205 | 194 |
| Initial CV * self-control | 250 | 218 | 217 |
| Initial UV * self-control | 132 | 129 | 122 |
| Initial direction * Initial chosen value | 183 | 166 | 157 |

**table S5.**
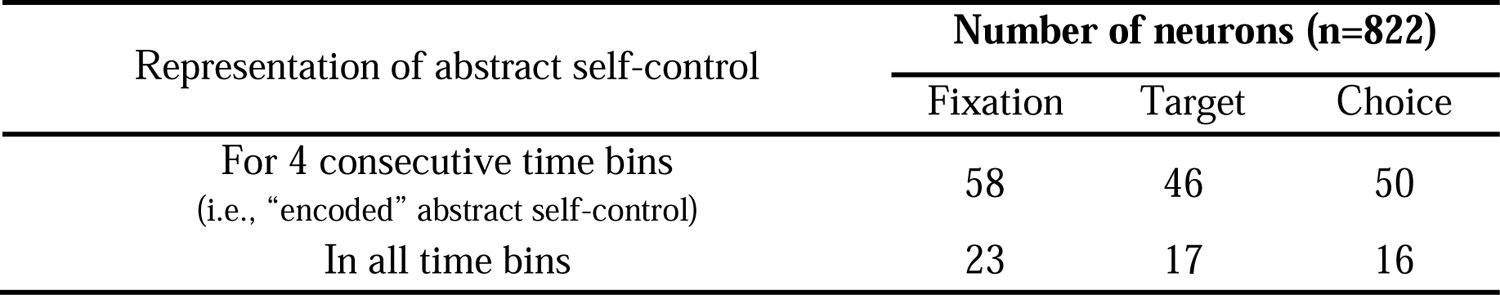
Instances of abstract self-control signals. Abstract self-control refers to encoding of self-control without an interaction with other task variables. First row counts the number of neurons that encoded abstract self-control (i.e., included abstract self-control in its best fitting model for at least 4 consecutive time bins). Note that this count can include neurons that encode an interaction with self-control in other time bins. Second row is the number of neurons that encodes abstract self-control in all time bins (i.e., self-control never has an interaction with other variables).

